# Environmental fluctuations do not select for increased variation or population-based resistance in *Escherichia coli*

**DOI:** 10.1101/021030

**Authors:** Shraddha Madhav Karve, Kanishka Tiwary, S Selveshwari, Sutirth Dey

**Affiliations:** *Population Biology Laboratory, Biology Division, Indian Institute of Science Education and Research-Pune*, *Dr. Homi Bhabha Road, Pune, Maharashtra 411008,* India.; *Present address: Advanced Center for Training, Education and Research in Cancer, Kharghar*, *Navi Mumbai, Maharashtra, 410210,* India.

**Keywords:** evolvability, standing variation, experimental evolution, antibiotic resistance, neutral space

## Abstract

Little is known about the mechanisms that enable organisms to cope with unpredictable environments. To address this issue, we used replicate populations of *Escherichia coli* selected under complex, randomly changing environments. Under four novel stresses that had no known correlation with the selection environments, individual cells of the selected populations had significantly lower lag and greater yield compared to the controls. More importantly, there were no outliers in terms of growth, thus ruling out the evolution of population-based resistance. We also assayed the standing phenotypic variation of the selected populations, in terms of their growth on 94 different substrates. Contrary to expectations, there was no increase in the standing variation of the selected populations, nor was there any significant divergence from the ancestors. This suggested that the greater fitness in novel environments is brought about by selection at the level of the individuals, which restricts the suite of traits that can potentially evolve through this mechanism. Given that day-to-day climatic variability of the world is rising, these results have potential public health implications. Our results also underline the need for a very different kind of theoretical approach to study the effects of fluctuating environments.

## 1 INTRODUCTION

The last few decades have witnessed a global increase in day-to-day climatic variability (Medvigy *et al.*, 2012). As a result of this, many organisms are now subjected to environmental changes at much shorter time scales than what they would have probably experienced for much of their evolutionary history. This has led to a number of empirical (reviewed in Hedrick, 2006, Kassen, 2002) and theoretical (Ishii *et al.*, 1989, Levins, 1968, Taddei *et al.*, 1997) studies, that seek to investigate the effects of environmental variability on the physiology (Hagemann, 2011) and evolution (Coffey *et al.*, 2011, Ketola *et al.*, 2013) of organisms. The primary insight that has emerged from these studies is that various aspects of the environmental heterogeneity, e.g. the number of components that constitute the environment (Barrett *et al.*, 2005, Cooper *et al.*, 2010), the speed with which the environment changes (Ancel, 1999, Cohan, 2005, Meyers *et al.*, 2005) or the predictability of environmental changes (Alto *et al.*, 2013, Hughes *et al.*, 2007) - can act singly, or in combinations with each other, to affect the evolutionary trajectory of populations. More interestingly, such fluctuations can lead to very different patterns of fitness in different test environments. For instance, in a recent study, when replicate populations of *E. coli* were subjected to fluctuating complex environments (random, stressful combinations of pH, salt and H_2_O_2_), the selected populations had no fitness advantage over the controls in stresses in which they were selected (i.e. pH or salt or H_2_O_2_ or combinations thereof) (Karve *et al.*, 2015). Yet, the same selected populations had significantly greater fitness in completely novel environments that had never been encountered by the bacteria before and had no known correlation with the stresses under which they had been selected (Karve *et al.*, 2015). Similar patterns of advantage under novel environments have been observed in other bacteria (Ketola *et al.*, 2013) and viruses (Turner *et al.*, 2000), when subjected to fluctuating selection pressure (although see Coffey *et al.*, 2011). These results are consistent with the general observation that disturbed habitats give rise to a large number of invasive species which, by definition, have fitness advantages in novel environments (Lee *et al.*, 2008 and references therein).

Unfortunately, in spite of a considerable corpus of theoretical predictions (Levins, 1968, Meyers *et al.*, 2005), there is little empirical work on the mechanisms that allow organisms to adapt to novel environments. Two major ways by which organisms can have greater fitness in novel environments are through an enhanced capacity to generate adaptive variations or by possessing larger amount of standing genetic variation. Although several organisms are known to respond to stress through increased mutation rate (Bjedov *et al.*, 2003) or enhanced phenotypic variation (Rohner *et al.*, 2013), it is not clear whether such traits can evolve due to exposure to environmental fluctuations. Recently, it has been shown that exposure to complex fluctuating environments do not lead to a significant change in mutation rates in *E. coli* (Karve *et al.*, 2015). However, it is hard to generalize on the issue as empirical studies on evolutionary effects of environmental fluctuations often do not investigate changes in mutation rates. The situation is not much different *w.r.t* the evolution of standing genetic variation under fluctuating environments. It has been shown *in vitro* that accumulated cryptic genetic variation in ribozymes can increase fitness in novel environments (Hayden *et al.*, 2011). However, chikungunya virus populations selected under fluctuating environments, show much less increase in genetic diversity compared to those raised in constant environments (Coffey *et al.*, 2011). Again, generalization of any kind is difficult, since we could not locate any other study that reports the changes in genetic variation in response to selection in fluctuating environments.

Apart from these variation-based mechanisms, there are a few other ways in which organisms can potentially deal with novel environments. The most well investigated of these seems to be phenotypic plasticity (Fusco *et al.*, 2010) which is expected to evolve when the environment changes faster than the life span of the organisms (Ancel, 1999, Meyers *et al.*, 2005). Another potential mechanism in this context might be an increase in broad-spectrum stress tolerance which is consistent with a recent finding that enhanced efflux activity evolves in *E. coli* in response to selection in fluctuating environments (Karve *et al.*, 2015). The third possible way to have greater fitness in novel environments is the evolution of population-based resistance, wherein a small fraction of individuals in the population synthesize some chemicals into the environment, which allows the entire population to become stress resistant (Lee *et al.*, 2010, Vega *et al.*, 2012). This kind of division of labor, in principle, can allow the population to become resistant to a wider spectrum of environments, thus enabling them to have greater fitness in a multitude of novel environments.

Here we investigate two of the above mentioned mechanisms for improved fitness in novel environments. We use replicate *E. coli* populations previously selected under unpredictable, complex environmental fluctuations for ~170 generations (Karve *et al.*, 2015). We test whether the phenotypic variation of the selected populations, in terms of usage of 94 substrates, have sufficiently diverged from the controls or not. We also count the number of progenies produced by individual bacterial cells, to ascertain whether population-based resistance has evolved in our selected populations. We find that our selected populations retain the fitness advantage even at the level of individual cells. However, there was no evidence of evolution of either increased phenotypic diversity or population-based resistance. Thus we can say that environmental fluctuations do not lead to increased variation, at least in the short time scale.

## 2 MATERIALS AND METHODS

### 2.1 Selection under constant and fluctuating environments

In this study, we used three replicate populations (henceforth F populations) of *E. coli* (strain NCIM 5547) that had been previously selected in unpredictably fluctuating, complex stressful environments. During the process of selection, the F populations were subjected to stressful combinations of salt, hydrogen peroxide and pH that changed unpredictably every 24 hrs. We also maintained corresponding controls (henceforth S populations) in the form of three replicate *E. coli* populations that were passaged in Nutrient Broth (see S1 for composition). After 30 days of selection (~170 generations, see S1 for calculations) these S and F populations were stored as glycerol stocks at -80^0^C. The details of the maintenance regime for both the F and the S populations have been mentioned elsewhere (Karve *et al.*, 2015).

### 2.2 Fitness of the individual bacteria in novel environments

To estimate the fitness of individual bacterium and characterize the possible heterogeneity within a population, we employed a slide-based observation technique (Lele *et al.*, 2011). Pilot studies were conducted to determine the sub-lethal concentrations for the four novel environments when the bacteria were grown on slides (see Table S2 for concentrations). The identity of these novel environments were chosen such that there are no known correlations between the mechanism of stress resistance to them and the three stresses used in the fluctuating selection (Karve *et al.*, 2015). Glycerol stock for S or F population was revived overnight in 50 ml Nutrient Broth. This revived culture was used to flood the slide layered with nutrient agar (see S1 for composition) containing one of the novel environment. After the broth had dried off (~ 30 minutes at room temperature under aseptic conditions), the agar surface was covered with a cover slip, excess agar outside the cover slip was removed with the help of a scalpel, and the sides were sealed with the mounting medium DPX (Di-n-butyl phthalate in xylene). The slide was then placed on the stage of a microscope (Primo Star™, Zeiss, Jena, Germany) which in turn was placed at 37^0^ C throughout the observation time.

A suitable field containing 6 to 20 single, well-spaced cells was focused under 100X magnification. For each cell in the field of view, we manually scored the time taken by the cell and its progenies to divide over a period of 240 minutes from the preparation of the slide i.e. from the time when broth was poured on the agar slide. Two trials were conducted for every replicate population of S and F in every novel environment (2 × 6 × 4 = 48 trials). The yield of each cell was estimated as the number of progenies produced by the cell at the end of 240 minutes. We also measured the ‘lag’ as the time taken for the first division. Since the cells were not synchronized, the lag estimate is likely to be associated with some amount of error. However, there is no reason to believe that this would affect S and F populations differentially. Moreover, since we measured substantial number of cells per population, such errors arising due to lack of synchronicity should be further ameliorated.

The yield and lag data were analyzed separately using mixed model ANOVA with novel assay environment (4 levels: Cobalt, Zinc, Norfloxacin and Streptomycin) and selection (2 levels: S and F) as fixed factors and replication (3 levels, nested within selection) and trial (2 levels, nested in assay environment × selection × replication) as random factors.

We also performed the individual mixed model ANOVAs for each of the novel assay environments. For this set of analysis, selection (2 levels: S and F) was treated as a fixed factor and replication (3 levels, nested within selection) and trial (2 levels, nested in selection × replication) as random factors. For the control of family-wise error rate, we used sequential Holm-Šidàk correction of the *p* values (Abdi, 2010). All ANOVAs in this study were performed on STATISTICA v5 (Statsoft Inc., Tulsa, OK, USA).

To estimate the effect size of the differences between the means, we computed Cohen’s *d* (Cohen, 1988) using the freeware Effect Size Generator (Devilly, 2004). The effect sizes were interpreted as small, medium and large for 0.2 < *d* < 0.5, 0.5 < *d* < 0.8 and *d* > 0.8 respectively (Cohen, 1988).

### 2.3 Population based resistance in novel environments

Population-based resistance occurs when a small fraction of the individuals synthesize a chemical which is then available to the other individuals of the population. However, as in our assay for individual-level fitness, when the bacteria are immobilized over an agar surface at extremely low densities for short durations, exchange of such chemicals become almost impossible. Thus, only those bacteria can resist the stresses which are able to synthesize the stress-fighting chemical on their own. If such bacteria are an extremely small fraction of the population, then they are expected to show up as outliers in the growth rate assay (see Discussion for further elaboration).

Most formal tests of outlier detection assume the underlying data to be normally distributed (Barnett *et al.*, 1978). Since our yield data did not meet this assumption, we used plots of the cumulative yield percentage to check for outliers. For this, we computed the percentage contribution of each parental bacteria to the final yield, arranged the values from both trials in ascending order and plotted the cumulative percentage yield against the percentage of the parental cells. In this plot, any cell(s) with disproportionate contribution to the overall yield can be easily identified by a sharp upward inflection towards the right of the graph.

### 2.4 Assay for phenotypic variation

We assayed the phenotypic variation in the population using GEN III MicroPlate™ (Catalog no. 1030 Biolog, Hayward, CA, USA). Each of these plates contains 94 separate substrates of which 71 can be utilized as carbon sources while 23 can act as growth inhibitors. The presence or absence of growth is indicated with the help of tetrazolium redox dye where intensity of purple color is proportional to the amount of growth.

From each of the F and S populations, we obtained 8 clones by streaking the glycerol stock on a Nutrient Agar plate and incubating overnight at 37^0^C. Thus a total of 48 clones were isolated over the three S and three F populations. Every clone was then characterized for the 94 different phenotypes on the Biolog plate using standard protocol (for detailed methods, see S3). An ancestral clone was processed in the same way to obtain the ancestral phenotypic profile.

Following a previous study (Cooper *et al.*, 2000), we measured absorbance of the plates at 590 nm using a microplate reader (SynergyHT BioTek, Winooski, VT, USA). For the 23 wells with inhibitory compounds, considering the recommendations of the product manual, we scored optical densities that were 50% or more of the corresponding positive control as 1 (i.e. no inhibition) and others as 0 (inhibition). Similarly, for the 71 wells with substrate utilization test, optical density that was ≥200% of the corresponding negative control was scored as 1 (i.e. utilized) while the others were scored as 0 (i.e. not-utilized). These binary scores were then used to determine standing phenotypic variation as well as the differences from the ancestral phenotypic profile. 33 phenotypes showed no variation in S and F (i.e. all individuals in S and F were either 0 or 1) and were ignored. For estimating standing phenotypic variation over the remaining 61 phenotypes, we computed the sum score of every replicate population over the eight clones. These values, ranging from 0 to 8, denote the variation within every population for that phenotype. It should be noted here that in some of the 94 substrates, absence of growth (i.e. 0) was the dominant phenotype while for the other substrates, the presence of growth (i.e. 1) was the dominant one. We were not interested in the qualitative nature of the phenotype (1 or 0) and wanted to analyze the variation over the entire set of 94 phenotypes. Therefore, we mapped phenotypic variation values of 5, 6, 7 and 8 to 3, 2, 1 and 0 respectively. In other words, a population in which three clones showed no growth (i.e. 3 zero values) and five clones showed growth (i.e. 5 values of 1), was deemed to have the same phenotypic variation for a given phenotype as a population which had five non-growers and three growers for a different phenotype. These mappings work only across phenotypes and fail if there are differences between the three replicates of S or F for the same phenotype. However, only three such cases were found in S populations and none at all in the F populations. The interpretations of our statistical analysis did not change with or without these points and hence we have retained these three data points. The phenotypic variations were then analyzed by a two way ANOVA with phenotype (61 levels) and selection (2 levels: S and F) as fixed factors. We also analyzed this data using the non-parametric Friedman test (Sokal *et al.*, 1995) after averaging over the three replicates populations for S and F. The inferences from both kinds of statistical tests were the same. Therefore we present the parametric analysis (i.e. 2-way ANOVA) here and discuss the non-parametric analysis, and its strengths and weaknesses, in the SOM (S7).

For estimating the phenotypic divergence from the ancestor, we recorded the number of clones displaying phenotype that was different from the ancestral one, for all the F and S populations. The number of differences for each phenotype was then analyzed using a two way ANOVA with phenotype (61 levels for Phenotypes) and selection (2 levels: S and F) as fixed factors.

## 3 RESULTS

### 3.1 Fitness of individual bacterial cells

When pooled over all the novel environments, individuals from F populations displayed significantly lower lag time (Fig 1A) and higher yield (Fig 1B) than individuals from S populations, with medium and high effect sizes respectively (Table 1). There was a significant effect of the novel environment in both cases (*F*_3, 12_ = 66.75, *p* < 0.001 for lag and *F*_3, 12_ = 88.93, *p* < 0.001 for yield) indicating that the difference in the fitness varies across different novel environments. This is intuitive as all the environments are not expected to affect fitness similarly. When analyzed separately for each novel environment, F populations had significantly and marginally significantly lower lag time in cobalt and streptomycin respectively (Table S4, Fig 1A) and significantly higher yield in cobalt, streptomycin and norfloxacin (Table S4, Fig 1B). The effect sizes were large in all these cases (Table S4). It is important to note that in all the four novel environments, F populations showed lower lag time and higher yield compared to S populations. The trends were consistent even when the two trials were analyzed separately (see SOM S5 for rationale and the results) which underlines the reproducibility of our results. However, these quantitative differences were not accompanied by any differences in cell size (F_1,4_ = 1.56, *p* = 0.279, *d* = 0.19) or shape (see SOM S6 for details).

**Table 1:** Summary of the main effects of selection in the pooled ANOVAs. Effect size was measured as Cohen’s *d* statistic and interpreted as small, medium and large for 0.2 < *d* < 0.5, 0.5 < *d* < 0.8 and *d* > 0.8 respectively.

| Assay | Means | | ANOVA F<br>(df effect, df error) | ANOVA P | Effect size<br>$\pm 95\% \text{CI}$ | Interpretation |
| --- | --- | --- | --- | --- | --- | --- |
|  | S | F |  |  |  |  |
| Lag | 147.66 | 118.69 | 19.20 (1,4) | 0.012 | 0.50±0.18 | Medium |
| Yield | 3.89 | 6.26 | 92.94 (1,4) | 0.0006 | 0.94±0.19 | Large |
| Phenotypic variation | 0.61 | 0.51 | 1.51 (1,244) | 0.22 | 0.11±0.2 | Small |
| Phenotypic divergence from ancestor | 0.97 | 0.79 | 3.98 (1,244) | 0.047 | 0.1±0.2 | Small |

**Fig 1:**
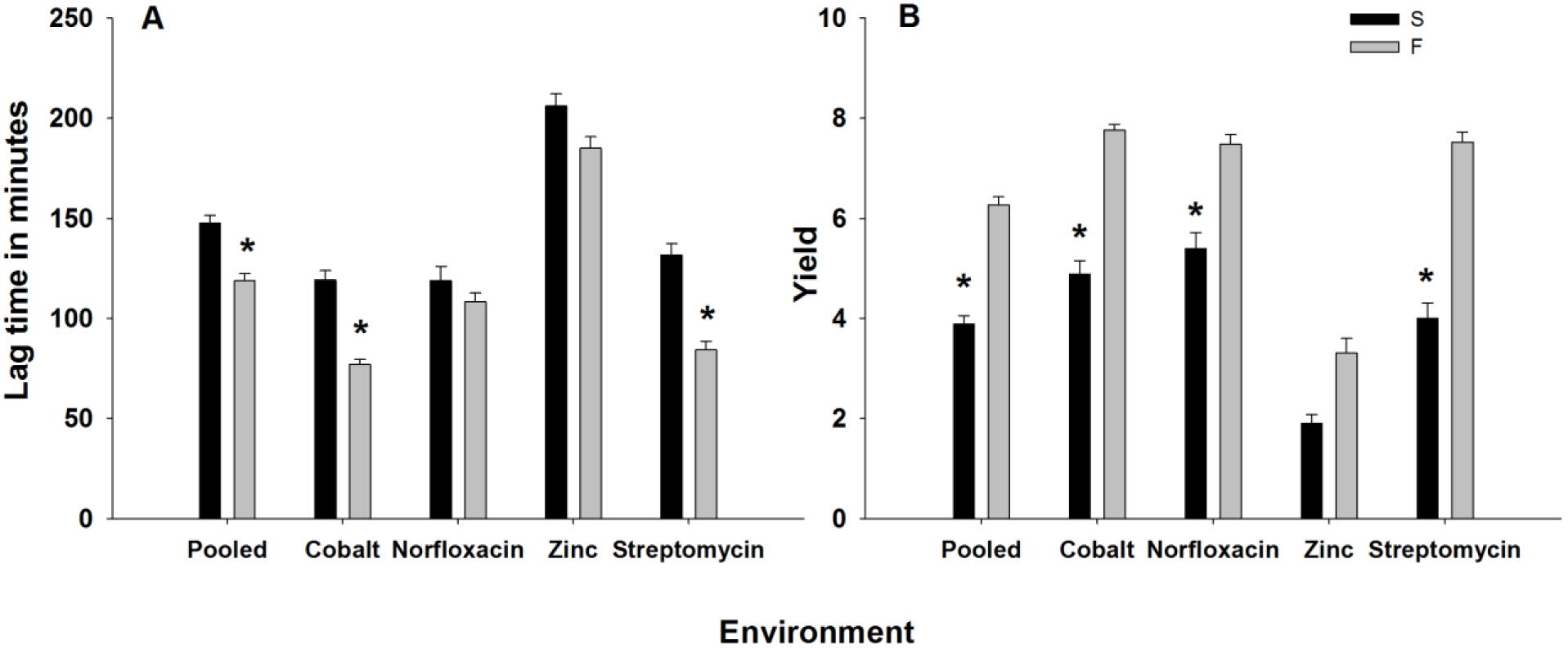
Fitness of individual bacterial cells. **A.** Mean (±SE) lag time is significantly lower for F populations than S populations when pooled over four novel environments. When compared separately for each novel environment, F populations show significantly lower lag time in cobalt and streptomycin and similar lag time in norfloxacin and zinc. **B.** Mean (±SE) yield is significantly higher for F populations than S populations when pooled over four novel environments. When compared separately for each novel environment, F populations show significantly higher yield for cobalt, norfloxacin and streptomycin and similar yield for zinc. * denotes *p* < 0.05 (after Holm-Šidàk correction in the case of comparisons under individual environments).

Overall, these results demonstrate the growth advantage for individuals of F populations in the four novel environments, corroborating the population level outcomes observed in an earlier study (Karve *et al.*, 2015).

### 3.2 Population-based resistance

Inspection of the data suggested that there were no individual cells whose progeny contributed disproportionately to the final population size. This can also be seen from the plot of the cumulative percentage yield of the cells, where the F populations showed a linear trend in three out of the four novel environments (Fig 2). Only in zinc, there was a small departure from the linearity (Fig 2D). However, even then ~20% of the cells contributing to ~40–60% of the observed yield and hence, there was nothing to suggest the presence of a small number of outliers that contributed disproportionately to the growth. Interestingly, zinc was the only novel environment where F populations did not display a fitness advantage in terms of yield or lag (see Discussion), thus ruling out the possibility of a few individuals conferring fitness advantage to the entire population.

**Fig 2:**
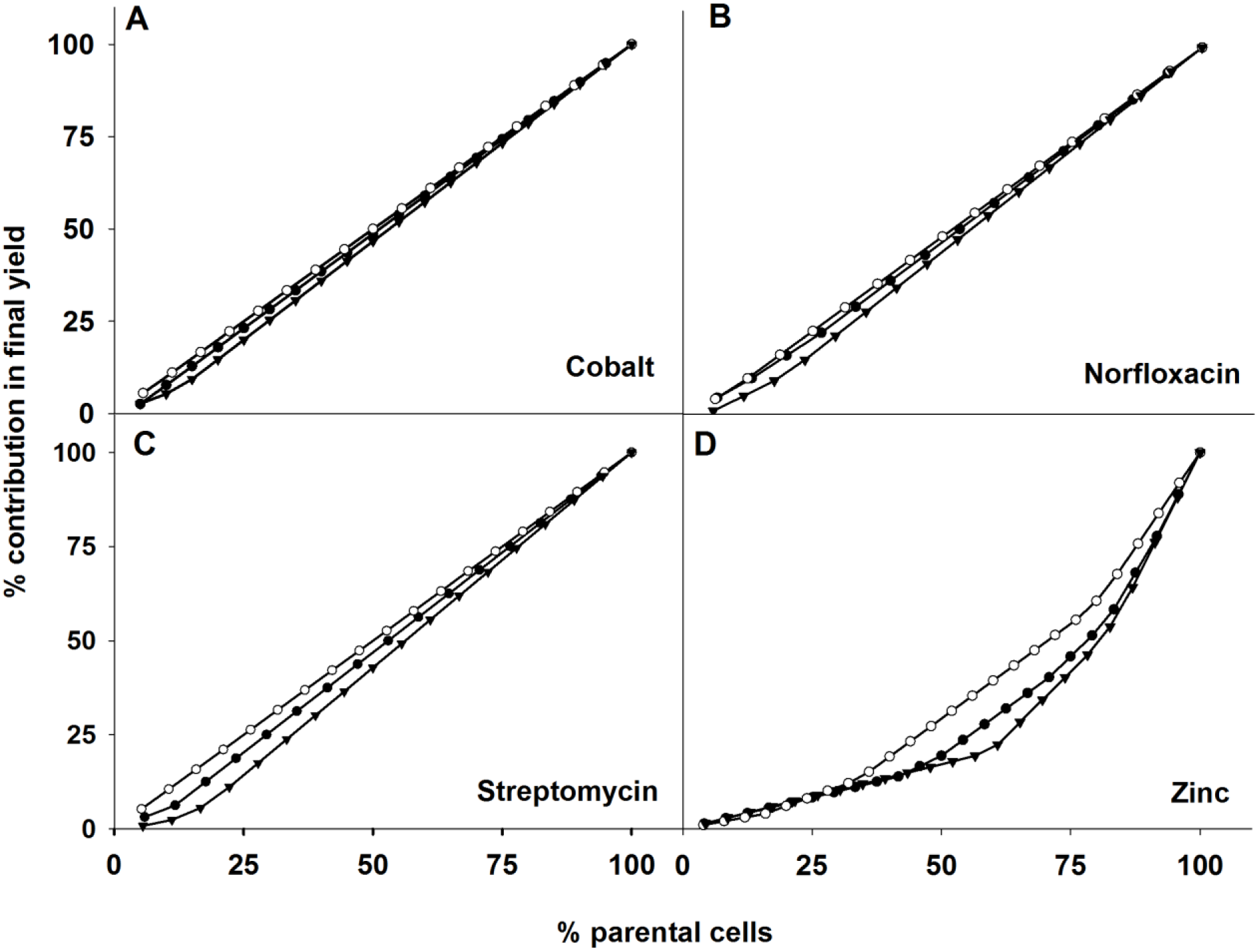
Population-based resistance in F populations. The cumulative percentage contribution of parental bacteria to the final yield is plotted for three replicate F populations in four novel environments. Each line in a figure stands for a replicate population of F. **A.** Cobalt, **B.** Norfloxacin, **C.** Streptomycin, **D.** Zinc. In this kind of a graph, the presence of outliers is detected as a sharp inflection towards the right, which was not observed. This indicates that no individual cells contributed disproportionately to the total yield.

### 3.3 Phenotypic Variation

For 33 out of the 94 substrates tested, no variation was found i.e. all the 48 clones of S and F gave the same phenotype. In the remaining 61 substrates, at least 1 out of the 48 clones (8 clones each for three S and three F populations) gave a different phenotype. ANOVA on the phenotypic distances showed a significant main effect of phenotype (*F*_60, 244_ = 3.69, *p* << 0.001) suggesting some phenotypes harbored more variation than others. This is intuitive as one does not expect similar number of variation for 61 traits over six populations. However, more crucially, there was no significant difference for the phenotypic variation across S and F populations, with a low effect size for the difference (Table 1 Fig 3A). Thus, we conclude that there was no evidence of an increased phenotypic variation in the F populations.

**Fig 3:**
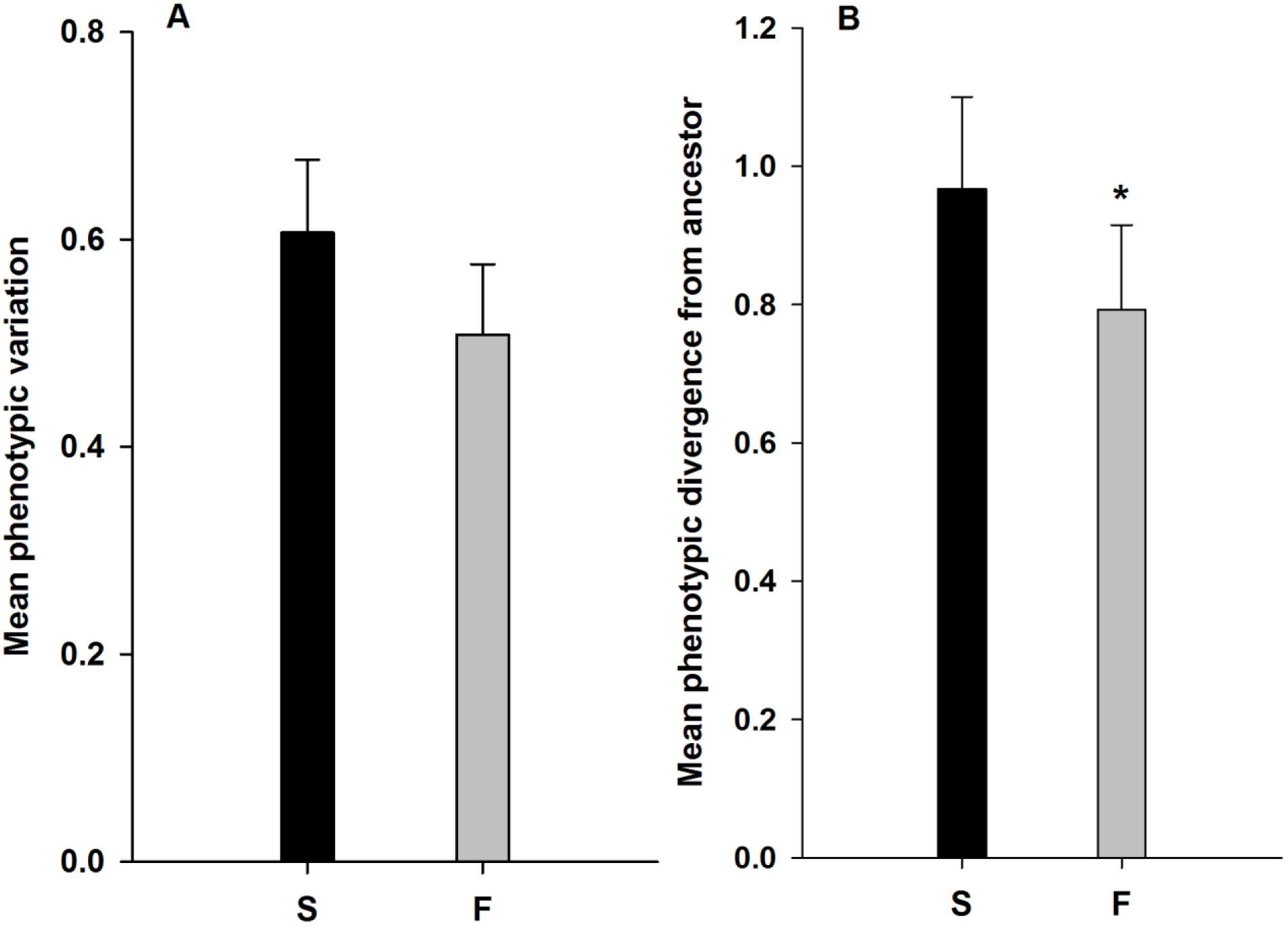
Phenotypic variation and divergence from ancestors. **A.** Mean (±SE) phenotypic variation for S and F populations. **B.** Mean (±SE) phenotypic divergence from the ancestors. The S populations show slightly higher variation and divergence albeit with small effect sizes. * denotes *p* < 0.05.

61 out of 94 phenotypes (not the same 61 as above though) showed at least one clone that was phenotypically different from the ancestor. Although, averaged over the 61 phenotypes, the S populations showed greater divergence which was marginally statistically significant (Fig 3B) the corresponding effect size of the difference was low (Table 1). More crucially, there was no phenotype for which all S or F populations were different from the ancestor. Barring two cases, no consistent pattern was observed in terms of acquiring or losing a phenotypic trait. 43 out of all the 48 clones tested acquired the ability to utilize methyl pyruvate while 39 became capable of utilizing β-methyl-D-glucoside. Although prior studies indicate that the ability to catabolize methyl pyruvate (Timonen *et al.*, 1998) and β-methyl-D-glucoside (Perkins *et al.*, 2008) often evolves under different kinds of stresses, the reason for the same remains unknown. Since, both S and F populations acquired the ability to utilize these compounds it is possible that there is some fitness advantage of these two phenotypes in nutrient broth. Crucially, there were no clear patterns in terms of phenotypic divergence from the ancestor, indicating that the variation accumulated is likely to be either neutral or have very weak effect on fitness. Apart from one replicate population of S in which all the individuals tested had lost the ability to utilize D-raffinose and pectin as a carbon source, there was not a single population in S or F in which all eight individuals had diverged from the ancestor. This suggests that the observed phenotypic variation is unlikely to be a result of a strong and /or directional selection pressure on one of the phenotypes. The divergence from ancestral phenotype varied significantly across different phenotypes (*F*_60, 244_ = 18.82, *p* << 0.001) with a significant interaction with selection (*F*_60, 244_ = 2.98, *p* << 0.001). Both these results are intuitive since one neither expects similar levels of divergence over 61 substrates nor similar patterns of divergence in S and F populations.

## 4 DISCUSSIONS

### 4.1 On measurement of fitness

Most experimental evolution studies in microbes measure fitness in either of two ways. The first is a measure of fitness in terms of growth rate or yield (Ketola *et al.*, 2013, Holder *et al.*, 2001). The second involves measuring competitive fitness by mixing the evolved strains with the ancestors and scoring their relative densities after a period of growth (Travisano *et al.*, 1995, Silander *et al.*, 2007). It is sometimes argued that the second method is more preferable as it also includes a measure of the competitive ability and hence gives an estimate of the magnitude of adaptation that has occurred in the selected populations over the course of the experiment (Kassen, 2014: page 16).

By definition, measuring competitive fitness equates evolutionary change with change in competitive ability and thus equates evolution with the ability of one genotype to replace another. It is, therefore, a narrow definition of fitness in the context of a correspondingly narrow and strict concept of evolution. However, the present study employs a broader notion of evolution as change through time within a species (Losos, 2013) and fitness as a measure of number of offspring in a given unit of time (i.e. yield) or any trait that affects that number (i.e. lag time). This is because we intuitively find no reason to expect that exposure to randomly fluctuating environments would lead to a change in competitive ability. Furthermore, we explicitly aimed to study fitness at the level of individual bacterium, which also enabled us to investigate phenomenon like population-based resistance. The notion of competitive fitness is not congruous with this aim and hence is not used here. To summarize, our concept of fitness is similar to the usage of Ketola *et al.* (2013) and Holder *et al.* (2001) and may or may not correspond with a change in competitive fitness as used in some other studies (Travisano *et al.*, 1995, Silander *et al.*, 2007).

### 4.2 Higher fitness of individuals in novel environments

For all the four novel environments, the lag times were lower for F populations and yields were higher (Fig 1) although the differences were not statistically significant for each comparison (Table S4). This corroborates similar observations at the population level in a previous study (Karve *et al.*, 2015). Unlike the population level assays, the individual level assays were conducted under an anaerobic environment. Though this could have affected the growth of the cells, the effect can be assumed to be similar across S and F individuals, and thus do not affect the conclusions of this study.

Increased fitness in multiple novel environments can come about in at least two major ways: an increased rate of generating new variation or the existence of larger amount of standing variation. If the first case were true, then one would not expect the progenies of all individuals of the F populations to acquire the favorable mutations at the same time in a novel environment. If the F populations had increased standing variation which was contributing to their enhanced fitness under novel environments, then again one would expect that most individuals would fail to grow and the progenies of only few individuals would primarily contribute to the final population size. However, we found no outliers in terms of contributions to the final size of the population (Fig 2) which suggests that whatever the mechanism that had evolved, was benefitting all the existing F individuals similarly. This observation does not fit with either increased rate of generation of variation or increased standing genetic variation.

### 4.3 No evidence for population-based resistance

When the magnitude and direction of selection fluctuates continuously, traits that are favorable under one set of conditions, might become neutral or even deleterious when the environment changes. This can lead to a scenario where a population is continuously changing with each shift in environment, without really evolving to greater fitness. One way by which a population can escape such a stasis is through the evolution of cooperation which allows subsets of the population to specialize in countering particular stresses and then confer resistance to the population as a whole (West *et al.*, 2007). For example, it has been shown that in populations of the bacteria *Pseudomonas aeruginosa,* the proportion of individuals that synthesize the iron-scavenging siderophore pyoverdin, changes based on the kind of competition and genetic relatedness (Griffin *et al.*, 2004). Similarly, when *E. coli* populations are challenged with antibiotics, a very small percentage (0.1 - 1%) of the individuals secret excess amounts of indole to the external environment, which then allows the entire population to become antibiotic resistant (Lee *et al.*, 2010). Since only a small fraction of the population needs to evolve the resistant mechanism for a given stress, in principle, this mechanism allows different subsets of the population to evolve resistance to different stresses. This should increase the population level variation in terms of the ability to resist diverse stresses, and hence increase fitness in different kinds of novel environments. Given that antibiotics were among the novel environments that we studied, population-based resistance was a possible explanation for the fitness advantages of F populations. Our assay for individual fitness was expected to detect the resistant subset as outliers with exceptionally high yield. This is because immobilization of cells at extremely low density over an agar surface limits the diffusion of extracellular metabolites over long distances and only those cells which synthesize the resistant factors can resist the stresses. However, we did not find any outliers in terms of the yield and, except in the case of zinc, all the plots of cumulative yield were linear (Fig 2). Even in the case of zinc, where there was a slight departure from linearity, at the point of inflection, ~20% of the parents contribute to ~40–60% of the yield. Overall, the conclusions are unambiguous, the observed increase in yield of the F populations were not attributable to a small fraction of the population.

The above result could have arisen in at least two other ways. It was possible that the F populations do exhibit population-based resistance, but we had managed to sample only those bacteria that conferred resistance to the population. The chances of such an event happening are probably negligible since, as stated already, the fraction of bacteria that confer the population-wide resistance is typically very low (Lee *et al.*, 2010). As we had sampled around 12-40 bacteria out of ~2 × 10^8^(over two trials) for each F population, it is highly unlikely that only individuals with altruistic capacities were sampled. In fact, the second possibility was far more likely, namely that we had sampled only those bacteria that did not confer any resistance to the population. In principle, this could also explain the absence of outliers in the F populations in terms of overall yield. However, in that case, we could not have observed an increase in the yield when compared to the S populations. Since the F populations did show a significantly larger yield compared to S populations (Fig 1B), we conclude that whatever mechanism was responsible for it, was not present only in a small number of individuals.

There can be multiple, non-exclusive reasons for which population-based resistance failed to evolve in our F populations. Our F populations were sub-cultured every 24 hours with 1/50 of the existing population forming an inoculum for the next generation (Karve *et al.*, 2015). It is difficult for population-based resistance to evolve in such a system due to a high chance of losing the resistant cells (which are in very low frequency) during each sub-culture. Moreover, it is known that when the environment changes, the production of the chemical that benefits the whole population can be costly for the producer cell (Lee *et al.*, 2010). Thus, in our F populations, there could have been a strong selection against the resistant cells, each time the environment changed. Taken together, perhaps it is not surprising that population-based resistance did not evolve in our F populations.

### 4.4 Fluctuating selection does not increase standing variation

Populations with greater standing variation are expected to respond faster to selection pressures compared to those with increased mutation rates. This is because with standing variation, the population need not wait for a beneficial mutation and such mutations are typically at a slightly higher frequency than those that arise *de novo* after exposure to the selection pressure (Barrett *et al.*, 2008). Furthermore, theoretical studies show that fluctuating environments are expected to promote standing variation in the populations (Turelli *et al.*, 2004, Gillespie *et al.*, 1989). Taken together, the greater fitness of the F populations in novel environments can be potentially explained if such populations have greater standing variation. We note here that a larger standing variation does not automatically guaranty that a population would be better able to face novel environments, it merely increases its chances for the same. However, it is difficult to visualize how large standing variation can be maintained when the direction of selection is changing very often (Via *et al.*, 1987). One way out of this problem is contextual neutrality, i.e. the assumption that at least some genetic changes are neutral in some environments (thus escaping selection) but affect fitness in other environments (thus contributing to standing genetic variance)(Wagner, 2005b). Thus, a population with a greater “neutral space” (i.e. contextually neutral variation) would be expected to have greater fitness across novel environments (Wagner, 2005a). Although some studies have directly measured genetic diversity through quantification of the number of mutations present (Coffey *et al.*, 2011), it is hard to determine how much of that diversity is functionally relevant. This is because, practically speaking, it is difficult to ascertain from the sequence data, whether a particular genomic mutation is deleterious, neutral or contextually neutral. Therefore, we favored a direct measurement of the phenotypic variation in the populations, through their ability to grow on 94 different conditions on the Biolog GEN III MicroPlate^TM^ plate (Cooper, 2002). This way, we quantify those variations that can cause an observable change at the phenotypic level and hence, are functionally important.

Our results suggest that selection for unpredictable fluctuations did not increase the phenotypic variation in F populations. If anything, the mean phenotypic variation was slightly larger for the S populations (Fig 3A), although the difference was not statistically significant. This is consistent with a previous study on viruses demonstrating that genetic diversity (as measured by genomic mutations) is larger in populations that experience a steady environment as opposed to those facing fluctuating ones (Coffey *et al.*, 2011). Our results are also in sync with a previous observation that constant selection environments lead to increase in the genetic variance for fitness in novel environments (Travisano *et al.*, 1995). In terms of the phenotypic divergence from the ancestors, we found no consistent differences or reversal of phenotypes that were specific to the F or S populations (Fig 3B).

There might be several reasons for which phenotypic variation did not increase in the F populations. The duration of selection (~170 generations) might have been too less to lead to a significant divergence in terms of phenotypic variation. Moreover, the fact that the environment (and hence the selection pressure) changed every ~6 generations, might have caused a much stronger selection pressure that prevented maintenance of phenotypic variation. One way by which standing variation can be increased even in the face of changing environments, is through increased mutation rates (Ishii *et al.*, 1989). However, since the mutation rates of the F populations did not evolve to be significantly larger than the S populations (Karve *et al.*, 2015), this route was closed to the selected populations. It is important to note here that we only scored the presence or absence of phenotypes, a process that is biased towards catching large phenotypic differences. In principle, one can also think of variations which affect the rate at which the substrates are metabolized or the intensity of the effect of stress substrates on the bacterial cells. However, quantifying such effects would require replicate measurements at the level of single clones and increased number of replicate clones due to the inherent variation in the metabolic rates of the cells, and hence was beyond the scope of this work.

### 4.5 Conclusion

Bacterial populations exposed to randomly fluctuating environments evolve to have greater fitness in novel stresses (Karve *et al.*, 2015, Ketola *et al.*, 2013). However, this is not attributable to an increase in standing variation, nor evolution of population-based resistance, nor an increase in the rate of generation of variation through mutations (Karve *et al.*, 2015). This suggests that the greater fitness in novel environments is perhaps due to direct individual-level selection on broad-spectrum stress resistance traits like change in membrane structure (Viveiros *et al.*, 2007), multi-drug efflux pumps (Nikaido *et al.*, 2012) etc. This observation is consistent with a previous result that the efflux activity of the F populations had increased significantly (Karve *et al.*, 2015). If the evolved increase in fitness were due to mutations or standing genetic variations, then there are a large number of ways available for the bacteria to evolve. However, the number of individual-level broad-spectrum resistance mechanisms is relatively small and typically well-studied, which at least gives some hopes in terms of developing containment strategies against such mechanisms. Moreover, most theoretical studies on evolutionary effects of fluctuating environments seek to model changes in mutation rates and standing variation (Ishii *et al.*, 1989, Leigh, 1970, Taddei *et al.*, 1997). Our results suggest that such studies have perhaps failed to consider the critical mechanism that enables organisms to adapt to such situations in nature and a new class of theoretical modeling is needed to investigate this issue.

## Acknowledgements

We thank Milind Watve, Yannis Michalakis and an unknown reviewer for helpful discussions and Madhur Mangalam for help with the single-cell assay. SK was supported by a Senior Research Fellowship from Council of Scientific and Industrial Research, Govt. of India. This study was supported by a research grant from Department of Biotechnology, Government of India and internal funding from Indian Institute of Science Education and Research, Pune.

## Supporting information

**S1** – Composition of Nutrient Agar and Nutrient Broth and Calculation of generation time in the experiment

**Table S2** – Novel Environments used for estimating fitness at the individual level

**S2** – Protocol for using the Biolog plates

**Table S4** – Summary of the main effects of selection in the ANOVAs under individual environments.

S5 – Summary of the main effects of the ANOVA for pooled data as well as under individual environments, for separate analyses for two trials

S6 – Quantitative analysis of the cell size

S7 – Nonparametric statistics for phenotypic variation

## Supplementary online material

### S1 Composition of Nutrient Agar and Nutrient Broth

Composition of Nutrient Agar (NA):

| Ingredients | g/L |
| --- | --- |
| Peptic digest of animal tissue | 5.00 |
| Sodium chloride | 5.00 |
| Beef extract | 1.50 |
| Yeast extract | 1.50 |
| Agar | 20.00 |

Final pH (at 25^0^C) 7.4 ± 0.2

The composition for Nutrient Broth (NB) is the same as above except the absence of agar.

For the slide based technique, we used 12 g / L of Agar.

#### Calculation of generation time in the experiment

We computed the number of generations (*g*) using the standard expression in the microbiology literature (Bennett *et al.*, 1997)
g = log_2_ (N^F^ / N_0_)
where, N^F^ is the population density (measured as OD) at the end of the growth cycle and N_0_ is the population density at the start of the growth cycle. It seems intuitive that F populations would have undergone less number of generations than S populations over the span of 30 days, due to the fact that the former were experiencing stress. However, this was not found to be the case, as seen from the data below:

| **Population** | **No. of generations** |
| --- | --- |
| **SA** | 170.50 |
| **SB** | 170.51 |
| **SC** | 170.37 |
| **HA** | 170.62 |
| **HB** | 170.54 |
| **HC** | 170.59 |

This seeming anomaly is explained by the fact that at every passage, we transferred fixed volume of culture to the next environment. Thus, though S populations reached higher OD_600_ value at almost every passage as compared to F populations, they started the next generation with higher numbers as well. As a result, the S populations hit stationary phase earlier than the F populations and, in effect, spent similar number of generations over the experimental duration.

**Table S2.** Novel Environments used for estimating fitness at the individual level.

| **Assay Environment** | **Concentration** |
| --- | --- |
| Cobalt chloride | 28.5 mg% |
| Zinc sulfate | 120 mg% |
| Streptomycin | 0.0065mg% |
| Norfloxacin | 0.0032 mg% |

### S3 Protocol for using the Biolog plates

We used GEN III MicroPlate™ along with inoculating fluid A (IF-A) for estimating the phenotypic variation. Both plates and inoculating fluid were stored at 4^0^C and thawed at room temperature before use.

A part of glycerol stock was streaked on nutrient agar plate for every replicate population of S and F. The plates were incubated at 37^0^C overnight. 8 isolated clones of comparable sizes were selected for every population and inoculated into the separate inoculation fluid tube. The transmittance was in the range of 95% to 98% for the selected clones. 100 μl of this well mixed inoculation fluid was used to inoculate the GEN III plate. The plates were incubated at 37^0^C for 24 hours after which they were stored at 4^0^C for another day, during which time we measured optical density for all the 48 plates at 590 nm (Cooper *et al.*, 2000) using a microplate reader (SynergyHT Biotek, Winooski, VT, USA).

**Table S4.**
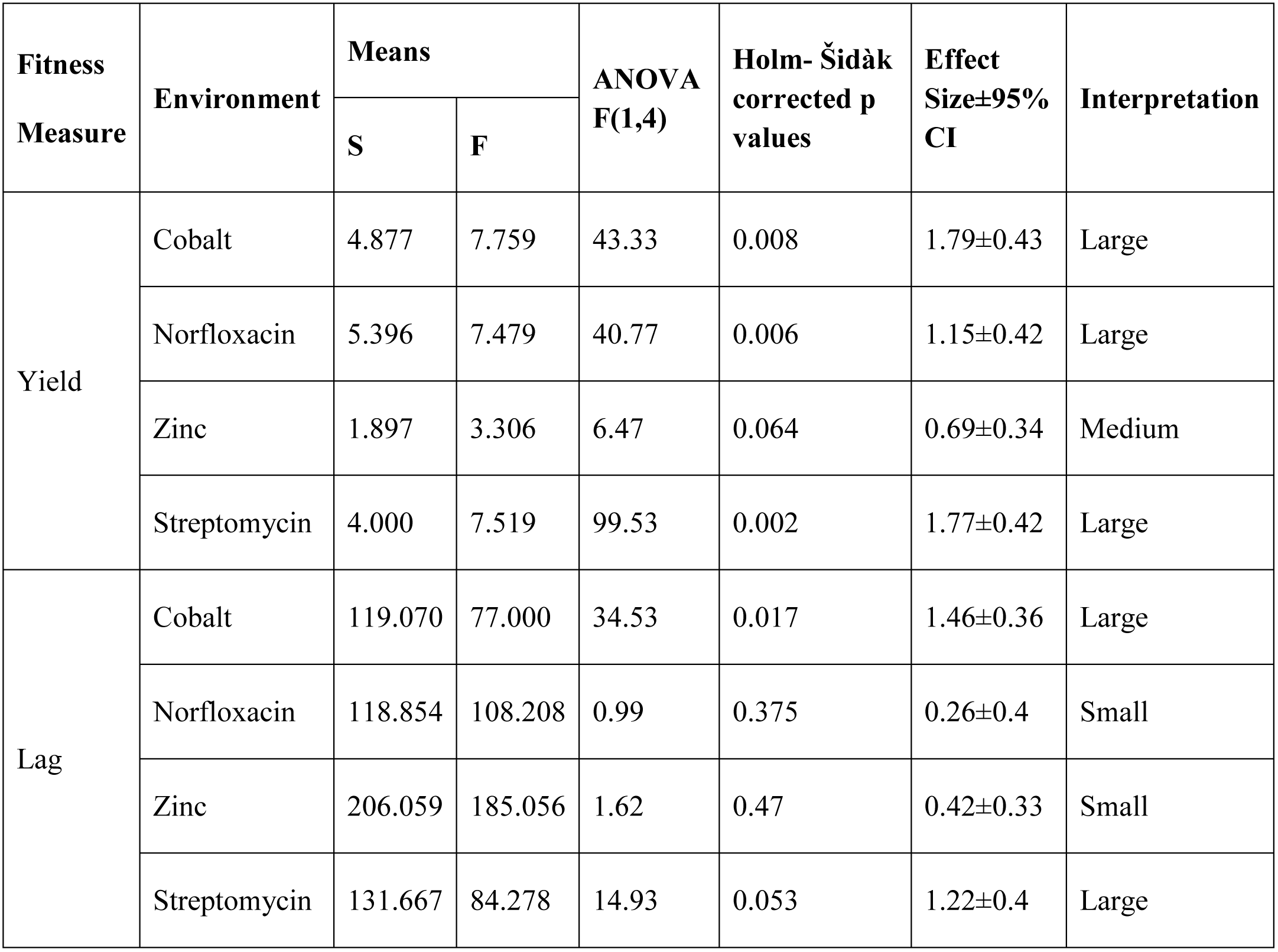
Summary of the main effects of selection in the ANOVAs under individual environments.

| Fitness Measure | Environment | Means |  | ANOVA F(1,4) | Holm- Šidák corrected p values | Effect Size±95% CI | Interpretation |
| --- | --- | --- | --- | --- | --- | --- | --- |
|  |  | S | F |  |  |  |  |
| Yield | Cobalt | 4.877 | 7.759 | 43.33 | 0.008 | 1.79±0.43 | Large |
|  | Norfloxacin | 5.396 | 7.479 | 40.77 | 0.006 | 1.15±0.42 | Large |
|  | Zinc | 1.897 | 3.306 | 6.47 | 0.064 | 0.69±0.34 | Medium |
|  | Streptomycin | 4.000 | 7.519 | 99.53 | 0.002 | 1.77±0.42 | Large |
| Lag | Cobalt | 119.070 | 77.000 | 34.53 | 0.017 | 1.46±0.36 | Large |
|  | Norfloxacin | 118.854 | 108.208 | 0.99 | 0.375 | 0.26±0.4 | Small |
|  | Zinc | 206.059 | 185.056 | 1.62 | 0.47 | 0.42±0.33 | Small |
|  | Streptomycin | 131.667 | 84.278 | 14.93 | 0.053 | 1.22±0.4 | Large |

This table shows yield and lag measurements for individual cells under four novel environments. Effect size was measured as Cohen’s *d* statistic and interpreted as small, medium and large for 0.2 < d < 0.5, 0.5 < d < 0.8 and d > 0.8 respectively.

### S5 Summary of the main effects of the ANOVA for pooled data as well as under individual environments, for separate analyses for two trials

We had not treated trials as repetitions at the level of the whole experiment because then we will have to conduct double the number of statistical tests than what we have done, which would have greatly inflated our overall error rates. Therefore, in the main manuscript, we treat trial as a random factor and analyze the data from the two trials together. However, as pointed out by a reviewer, the two trials can also be used to check for reproducibility of the results. Therefore, we have reanalyzed the individual cell data for two trials separately and find that the results are consistent with our earlier analysis.

Trial I

Main effect of selection in pooled data

| **Assay** | **Means** |  | **ANOVA F**<br>**(df effect,df error)** | **ANOVA**<br>**p** | **Effect**<br>**Size** | **Effect Size**<br>**Interpretation** |
| --- | --- | --- | --- | --- | --- | --- |
|  | **S** | **F** |  |  |  |  |
| Lag | 141.68 | 115.49 | 15.12(1,4) | 1.80E-02 | 0.5±0.26 | Medium |
| Yield | 3.80 | 6.33 | 43.03(1,4) | 0.003 | 1.04±0.28 | Large |

Main effect of selection in individual environments

| Assay | Environment | Means |  | ANOVA<br>F(1,4) | Holm-Sidak<br>Correction p values | Effect Size | Effect Size Interpretation |
| --- | --- | --- | --- | --- | --- | --- | --- |
| Lag |  | **S** | **F** |  |  |  |  |
|  | Cobalt | 123.13 | 77.57 | 11.36 | 0.108 | 1.3±0.57 | Large |
|  | Norfloxacin | 126.26 | 106.15 | 0.77 | 0.676 | 0.44±0.6 | Small |
|  | Zinc | 187.84 | 180.00 | 0.14 | 0.729 | 0.14±0.48 | Small |
|  | Streptomycin | 124.33 | 81.83 | 6.08 | 0.194 | 1±0.54 | Large |
| Yield | Cobalt | 4.07 | 7.71 | 44.55 | 0.01 | 2.6±0.7 | Large |
|  | Norfloxacin | 5.35 | 7.05 | 7.42 | 0.103 | 0.79±0.62 | Large |
|  | Zinc | 2.52 | 4.09 | 3.96 | 0.117 | 0.65±0.5 | Medium |
|  | Streptomycin | 3.67 | 7.17 | 35.8 | 0.012 | 1.61±0.58 | Large |

Trial II

Main effect of selection in pooled data

Trial II Main effect of selection in pooled data
| Assay | Means |  | ANOVA F | ANOVA p | Effect Size | Effect Size Interpretation |
| --- | --- | --- | --- | --- | --- | --- |
|  | **S** | **F** | **(df effect,df error)** |  |  |  |
| Lag | 153.39 | 121.72 | 7.27(1,4) | 0.054 | 0.54±0.26 | Medium |
| Yield | 3.97 | 6.20 | 30.96(1,4) | 0.005 | 0.85±0.27 | Large |

Main effect of selection in individual environments

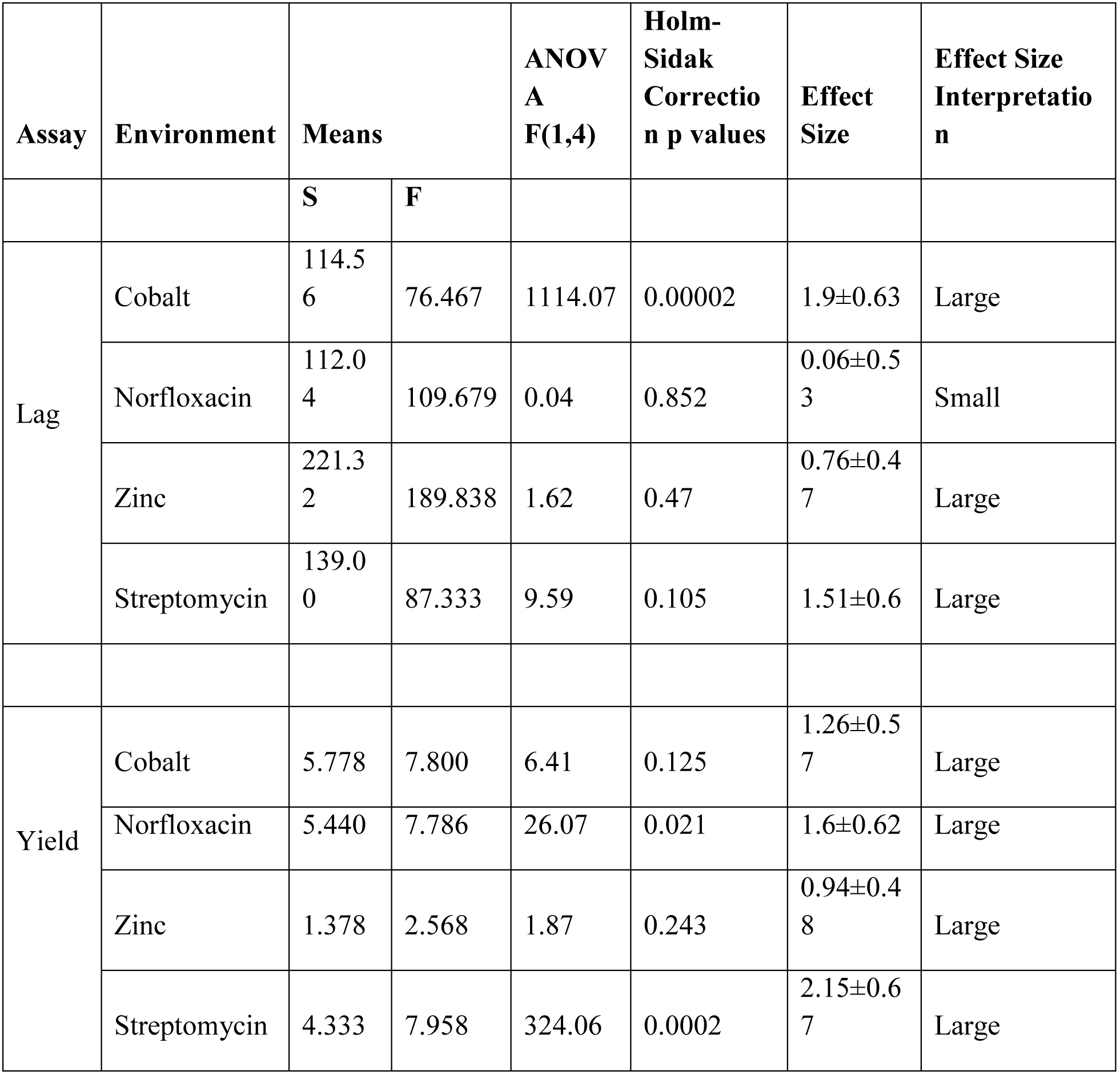

The trends were consistent even when the two trials are analyzed separately, which underlines the reproducibility of our results.

### S6 Quantitative analysis of the cell size

One of the ways in which the F populations might have evolved could have been by change in cell shape or cell size. To study this, we exposed individuals from all the replicate S and F populations to the same concentration of Norfloxacin as used in the study. We used ImageJ software (Schneider *et al.*, 2012) to measure the cell sizes right after the cells divided (to make sure that all cells are measured at the same age). Cell lengths were measured for first two divisions and data was analyzed using a 2-way ANOVA where selection (two levels – S and F) was a fixed factor and replicate (three levels) was a random factor nested in selection. The difference in cell sizes is non-significant with very small effect size (F_1,4_ = 1.56, *p* = 0.279, d = 0.19). We also provide representative images.

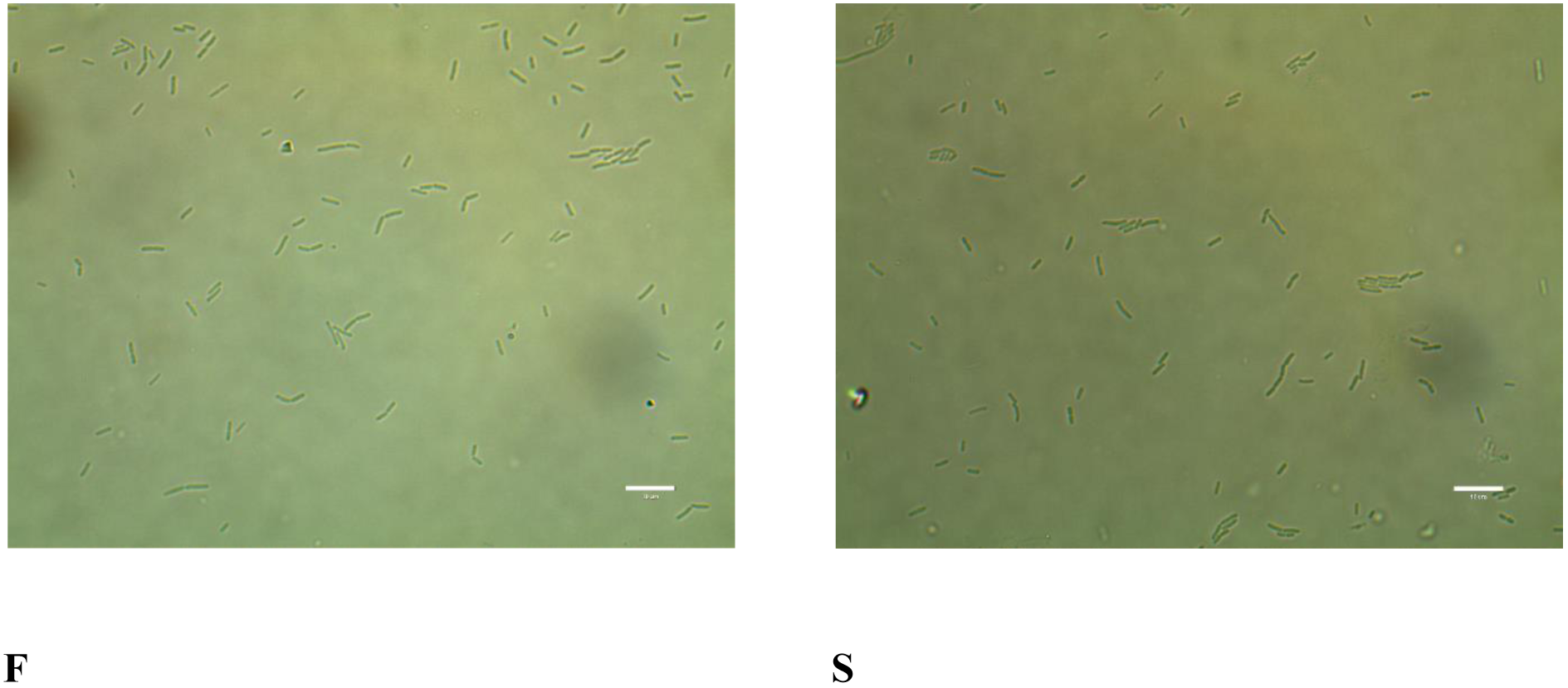

### S7 Nonparametric statistics for phenotypic variation

The discrete nature of the phenotypic variation scores violates the normality assumption of ANOVA (Zar, 1999). However, it is important to note that:

a. ANOVA is known to be robust to departures from normality, particularly when the numbers of observations in each cell are the same (Quinn *et al.*, 2002), which they are in this case.
b. The effect sizes for all comparisons have been computed and Cohen’s *d* does not depend on the nature of the underlying distribution. The interpretation from the effect sizes was the same as that of the parametric analysis: there were no major differences between the phenotypic variation of S and F.

In spite of this, as recommended by an editor, we conducted a non-parametric analysis of the data. The use of the non-parametric alternative of the two way ANOVA (Scheirer-Ray-Hare test) is controversial in the statistical literature (Dytham, 2011), and hence was not used here. Instead, we performed another commonly used nonparametric test, namely the Friedman test (Sokal *et al.*, 1995). Phenotypic distance for every environment was averaged over three replicate populations for S and F. This yielded 61 different values of distance each for S and F, every single value corresponding to one environment. This data was then analyzed using the Friedman test and we detected no significant difference in phenotypic variance between the S and F populations (*p* = 0.473) which is consistent with the results obtained from the 2-way ANOVA (*p* = 0.22; Table 1 of the manuscript). This is not surprising as non-parametric tests have considerably lower power than the corresponding parametric ones (Zar, 1999), and therefore any difference between means that is statistically non-significant in parametric tests will remain non-significant in non-parametric tests.

To summarize, whatever way we analyzed the phenotypic variance (ANOVA, effect size, Friedman test), the conclusions were the same: there was no significant difference between the phenotypic variance in S and F populations.

